# Tissue-specific cis-regulatory divergence implicates a fatty acid elongase necessary for inhibiting interspecies mating in *Drosophila*

**DOI:** 10.1101/344754

**Authors:** Peter A. Combs, Joshua J. Krupp, Neil M. Khosla, Dennis Bua, Dmitri A. Petrov, Joel D. Levine, Hunter B. Fraser

**Affiliations:** Department of Biology, Stanford University; Department of Biology, University of Toronto, Mississauga

## Abstract

Pheromones known as cuticular hydrocarbons are a major component of reproductive isolation in *Drosophila*. Individuals from morphologically similar sister species produce different sets of hydrocarbons that allow potential mates to identify them as a suitable partner. In order to explore the molecular mechanisms underlying speciation, we performed RNA-seq in F1 hybrids to measure tissue-specific cis-regulatory divergence between the sister species *D. simulans* and *D. sechellia*. By focusing on cis-regulatory changes specific to female oenocytes, we rapidly identified a small number of candidate genes. We found that one of these, the fatty acid elongase *eloF*, broadly affects both the complement of hydrocarbons present on *D. sechellia* females and the propensity of *D. simulans* males to mate with those females. In addition, knockdown of *eloF* in the more distantly related *D. melanogaster* led to a similar shift in hydrocarbons as well as lower interspecific mate discrimination by *D. simulans* males. Thus, cis-regulatory changes in *eloF* appear to be a major driver in the sexual isolation of *D. simulans* from multiple other species. More generally, our RNA-seq approach proved to be far more efficient than QTL mapping in identifying candidate genes; the same framework can be used to pinpoint cis-regulatory drivers of divergence in a wide range of traits differing between any interfertile species.

## Introduction

Reproductive isolation is a major component of speciation. Postzygotic incompatibilities leading to hybrid sterility or inviability (also known as Dobzhansky-Muller incompatibilities) have been especially well-studied, with several examples narrowed down to specific genes (Watanabe 1979; Sawamura *et al.* 1993; Phadnis *et al.* 2015). However when the distributions of related species overlap, rejection of interspecific partners may account for a much larger fraction of reproductive isolation (Coyne and Orr 1997; Quinn *et al.* 2000; Byrne and Rice 2006; Shahandeh *et al.* 2018). This preference for conspecific mates may be subject to strong selection (Noor 1995; Servedio and Noor 2003; Coyne and Orr 2004), since interspecific hybridization carries significant fitness costs, including potential inviability or sterility of offspring.

*Drosophila* has been a key model organism for the study of reproductive isolation, including the role of mate choice (Coyne and Orr 2004). Courtship in *Drosophila* is a highly stereotyped procedure, with multiple opportunities for both females and males to reject interspecific partners (Sokolowski 2001; Lasbleiz *et al.* 2006). This affords the opportunity for flies to reduce energy expenditure on reproductively fruitless partners. While female mate choice has been more heavily studied (Spieth 1952; Partridge 1980; Fowler and Partridge 1989; Greenspan and Ferveur 2000), there is a growing recognition that choice by males can also be an important factor (Byrne and Rice 2006; Edward and Chapman 2011; Pischedda *et al.* 2014; Shahandeh *et al.* 2018). In fact, male choice can be responsible for most reproductive isolation in some cases (Shahandeh *et al.* 2018). Beyond simply the opportunity cost of devoting time towards courting a heterospecific female, mating itself can be costly for males, with mated male *Drosophila* having reduced lifespans (Partridge and Farquhar 1981). Simulations have shown that male mate choice can reinforce speciation under when hybrids are less fit (Servedio 2007).

*D. simulans* and *D. sechellia* are two closely related sister species, separated by approximately 250 thousand years (Garrigan *et al.* 2012). The species are believed to have diverged in allopatry (Kliman *et al.* 2000), though currently their ranges overlap and hybrids can be found in the wild (Matute and Ayroles 2014). In laboratory conditions, *D. sechellia* males will readily mate with *D. simulans* females, producing sterile male and fertile female hybrid offspring, while the reciprocal cross is much more difficult (Lachaise *et al.* 1986). Male mate choice in these species—which accounts for over 70% of their reproductive isolation (Shahandeh *et al.* 2018)—is mediated by female cuticular hydrocarbons (CHCs), which are key molecules involved in species recognition that are produced primarily in specialized cells called oenocytes (Billeter *et al.* 2009).

In this study, we sought to identify the specific gene(s) responsible for CHC-mediated behavioral reproductive isolation in *D. simulans* and *D. sechellia*. Thus far, QTL mapping has been the primary method used to investigate this question. QTLs affecting CHCs have been mapped, but these contain many CHC-related genes (Coyne *et al.* 1994; Gleason *et al.* 2005; 2009), and fine-mapping has not been reported. As a complementary approach, we reasoned that genes responsible for major changes in CHCs may share three key characteristics: 1) Cis-regulatory divergence in female oenocytes; 2) Female-specific expression; and 3) Oenocyte-specific expression. Although these are certainly not required—for example, CHC divergence might occur via changes in protein-coding regions—any genes meeting all three criteria would be excellent candidates.

Cis-regulatory divergence can be measured genome-wide via high-throughput sequencing of cDNA (RNA-seq) in interspecific hybrids. Hybrids are required because comparisons between species involve a combination of both cis- and trans-acting changes; in contrast, measuring allele-specific expression (ASE) in F1 hybrids neatly controls for potential trans-acting changes, since each allele experiences the same trans-regulatory environment within the hybrid nuclei. Thus, differential expression of the two alleles in a hybrid can only be explained by cis-regulatory divergence.

To generate genome-wide data covering all three criteria listed above, we performed RNA-seq in *D. sechellia*/*simulans* hybrids. To measure female-specificity, we included samples from both male and female oenocytes, and to measure oenocyte-specificity, we included samples from male and female fat bodies (an adjacent non-CHC producing tissue; Lawrence and Johnston 1986). Using this approach, we identified three candidate genes for drivers of CHC differences between the species. Ablation of these genes pointed towards a major role of *eloF*, a fatty acid elongase, in the reproductive isolation of *D. simulans* from both *D. sechellia* as well as the more distantly related *D. melanogaster*.

## Results

### Allele-specific expression identifies fatty acid elongases as a major differentiator between *D. simulans* and *D. sechellia* female oenocytes

We first set out to identify genes with cis-regulatory divergence specific to female oenocytes. We mated *D. sechellia* males to *D. simulans* females and dissected both oenocytes and fat bodies from the progeny, pooling approximately 20 individuals from each sex (Figure 1A). Then, we extracted RNA and constructed RNA-seq libraries, which we sequenced to approximately 30 million reads per sample (Supplemental Table 1). We called allele-specific reads for each sample by aligning to a *D. simulans* reference sequence, and controlled for potential mapping bias by discarding any read that did not map to the same location if alleles were swapped *in silico* (van de Geijn *et al.* 2015). Despite the use of a *D. simulans* reference genome, we found a majority of reads were assigned to *D. sechellia* (Supplemental Table 1), possibly indicating low levels of non-hybrid *D. sechellia* samples. We estimated the significance of each gene’s allele specific expression (ASE) using a negative-binomial test (Love *et al.* 2014) for deviation from the average fraction of *D. sechellia* reads in a given sample.

**Figure 1:**
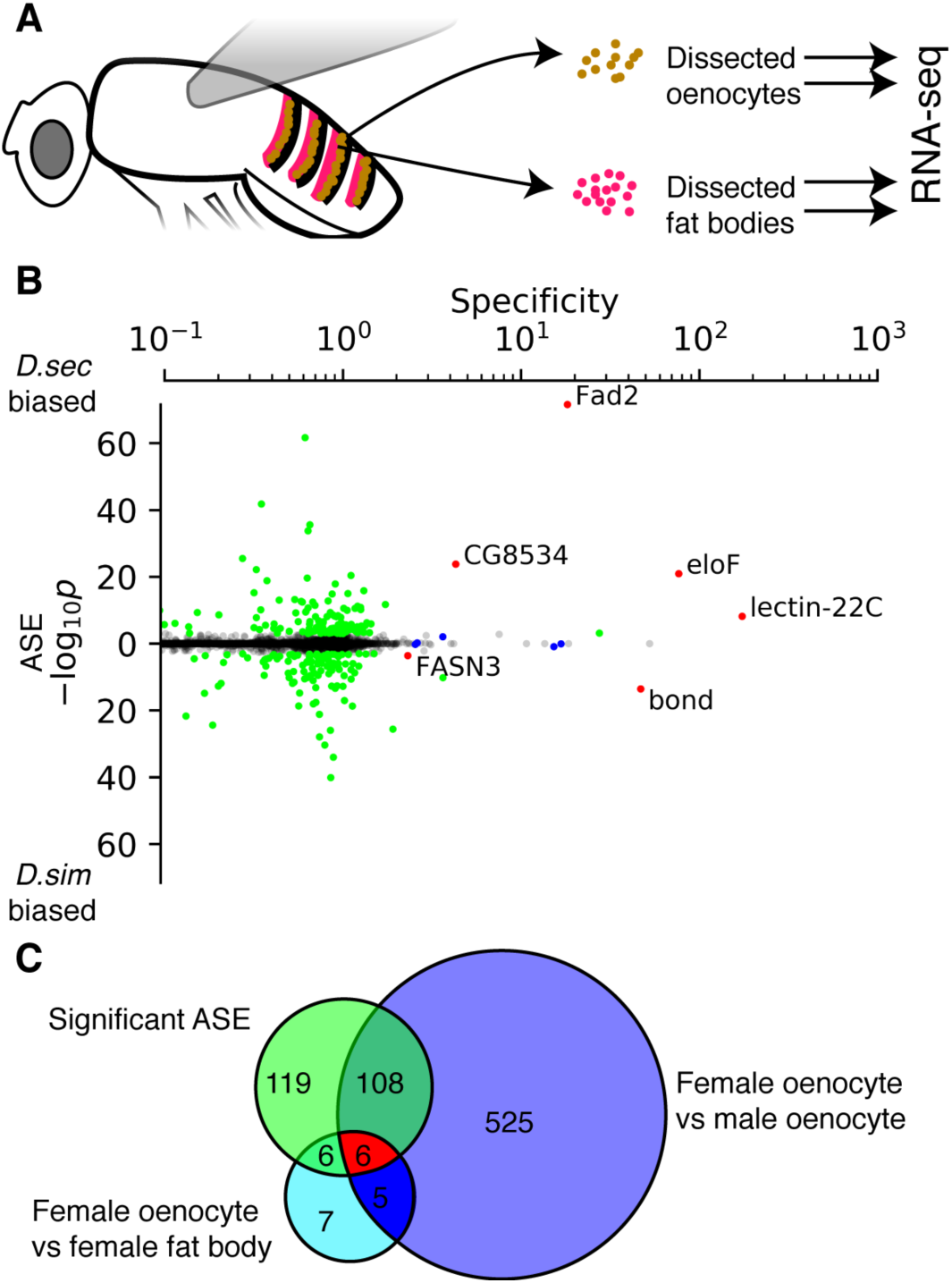
RNA-seq of oenocytes and fat bodies from hybrid *D. simulans × D. sechellia* flies reveals a strong cis-regulatory component of CHC production. **A)** We dissected oenocytes (blue dots) and fat bodies (green regions) from hybrid *D. simulans × D. sechellia* males and females and performed RNA-sequencing. **B)** Genes are plotted by specificity of expression to female oenocytes (x-axis; mean of female oenocyte expression divided by maximum expression in female fat bodies, male oenocytes, and female oenocytes) and allele-specific expression p-value (y-axis). Green dots indicate genes with significant ASE compared to the distribution of reads in the female oenocytes, blue dots indicate those that have significantly higher expression in female oenocytes compared to female fat bodies and male oenocytes, and red dots indicate genes with both tissue-specific and species-specific expression. **C)** Overlap of genes with ASE in female oenocytes (green circle), and differential expression in female oenocytes compared to other tissues (blue and cyan circles).

Even at a stringent cutoff, we identified 239 genes with significant (negative binomial q-value < 0.001) ASE in female oenocytes. This is not surprising, since various *Drosophila* interspecific hybrids have also yielded large numbers of genes with strong ASE. Of the 239 significant genes, 27 have been annotated with the Gene Ontology term “Fatty acid biosynthetic process” (GO:0006633) (Supplemental Table 3). Therefore we concluded that, even when combined with GO annotations, ASE in female oenocytes was insufficient to identify a manageable number of candidate genes involved in CHC differences and speciation.

We reasoned that in addition to ASE, genes important to female CHC differences between *D. simulans* and *D. sechellia* would likely be expressed specifically in female oenocytes (Figure 1B and C). To identify candidate genes, we looked for genes that had significantly higher expression in the female oenocytes compared to both male oenocytes and female fat bodies (Sleuth q-value<0.001 for both comparisons; (Pimentel *et al.* 2017)). Only six genes passed these cutoffs. Reassuringly, one of these was *desatF* (also known as *Fad2*), a fatty-acid desaturase which is known to be expressed in *D. sechellia* female oenocytes, but not in males or in *D. simulans* (Shirangi *et al.* 2009).

Among the six candidate genes, the only enriched molecular function Gene Ontology terms were related to “fatty acid elongase activity” (GO:0009922 and its parent GO terms), which describe the three genes *eloF, CG8534,* and *bond* (in all cases, we use the names of the *D. melanogaster* orthologs) (Boyle *et al.* 2004). All three of these have ELO family domains (Szafer-Glusman *et al.* 2008). Both *eloF* and *CG8534* were *D. sechellia-*biased, while *bond* was *D. simulans-*biased. We further detected a weak signal for *FASN3,* a putative acyl transferase (Table 1). No other gene that is both oenocyte- and species-specific in its expression has an annotated Gene Ontology term or protein domain that is clearly related to CHC production (Table 1).

**Table 1:** Genes with female oenocyte- and species-specific expression. Genes with significant tissue-specific (sleuth q-value <0.001 in comparisons both between the two female tissues, and between the two oenocyte samples) and species-specific expression (negative binomial p-value < .001). Specificity is the ratio of the mean expression in female oenocytes to the highest expression among male oenocytes, female fat bodies, and male fat bodies. Gene ontology (GO) terms are annotated molecular function terms (see Supplemental Table 2 for citations). GO terms without experimental evidence are in italics. Protein domains are InterPro annotated protein domains/motifs as listed on FlyBase v2017_06 (Finn *et al.* 2017; Gramates *et al.* 2017).

| *Gene* | *Female*<br>*Oenocyte*<br>*Specificity*<br>*(oenocyte/female*<br>*Sleuth q-value)* | *% D. sechellia*<br>*reads in female*<br>*oenocytes*<br>*(negative*<br>*binomial p-value)* | *GO term(s)* | *Protein Domain(s)* |
| --- | --- | --- | --- | --- |
| *eloF* | 76.6<br>(4.5e-9/2.3e-5) | 98.75%<br>(1.2e-21) | fatty acid elongase activity | ELO family |
| *Fad2* | 18.2<br>(4.5e-9/2.5e-5) | 95.5%<br>(2.5e-72) | *Catalysis of an oxidation-reduction (redox) reaction in which hydrogen or electrons are transferred from each of two donors ...* | Fatty acid desaturase type 1, conserved site; Fatty acid desaturase domain; Acyl-CoA desaturase |
| *CG8534* | 4.3<br>(8.2e-4/3.6e-5) | 93.75%<br>(1.6e-24) | *fatty acid elongase activity* | ELO family |
| *FASN3* | 2.31<br>(2.7e-4/3.8e-5) | 60.5%<br>(.00026) | *3-hydroxypalmitoyl-[acyl-carrier-protein] dehydratase activity; 3-oxoacyl-[acyl-carrier-protein] reductase (NADPH) activity; 3-oxoacyl-[acyl-carrier-protein] synthase activity; [acyl-carrier-protein] S-acetyltransferase activity; [acyl-carrier-protein] S-malonyltransferase activity; enoyl-[acyl-carrier-protein] reductase (NADPH, B-specific) activity; myristoyl-[acyl-carrier-protein] hydrolase activity; oleoyl-[acyl-carrier-protein] hydrolase activity; palmitoyl-[acyl-carrier-protein] hydrolase activity; phosphopantetheine binding* | Ketoacyl synthase (N-terminal, C-terminal, and C-terminal extension), Acyl transferase, Polyketide synthase, Alcohol dehydrogenase C-terminal |
| *lectin-22C* | 173.9<br>(3.0e-4/2.0e-4) | 85.75%<br>(6.7e-09) | *galactose binding* | C-type lectin-like/link domain superfamily |
| *bond* | 47.0<br>(7.8e-4/6.3e-5) | 25.5%<br>(2.8e-14) | fatty acid elongase activity | ELO family |

Compared to the female oenocytes, male oenocytes had a much weaker signal of ASE among genes with sex- and oenocyte-specific expression (Supplemental Figure 2). Given the overall weaker signal in male oenocytes, we chose to focus on changes in female CHC production that might drive speciation.

Male fat bodies had over 80 genes with tissue- and species-specific expression (Supplemental Figure 2A). Gene ontology analysis of these male fat body genes highlighted several significant GO terms, including “oxidation-reduction process” (p=2.9 × 10^−7^) and “catalytic activity” (p=6.7 × 10^−10^) (Boyle *et al.* 2004), but no candidate genes with obvious roles in pheromone production or mating activity were present. However, these genes may be useful for future studies of regulatory evolution in fat bodies, which could affect traits including metabolism and mating behavior (Lazareva *et al.* 2007).

### eloF has widespread effects on the hydrocarbon profile of *D. sechellia* and *D. melanogaster*

To explore the role of our candidate genes on CHC profiles of these species, we performed gas chromatography coupled to mass spectrometry (GCMS). Consistent with previous measurements of hydrocarbon profiles of *Drosophila*, we found that wildtype *D. simulans* has more short-chain hydrocarbons than *D. sechellia* (Figure 2A; (Jallon and David 1987)). In particular, *D. sechellia* has almost no 23-carbon CHCs, while the predominant *D. simulans* hydrocarbon is 7-tricosene, a 23-carbon monoene. Indeed, there was only one hydrocarbon shorter than 26 carbons with a greater representation in *D. sechellia* than *D. simulans,* the 25-carbon pentacosadiene (~2 fold higher in *D. sechellia)*. There were no CHCs longer than 26 carbons that were more abundant in *D. simulans* than *D. sechellia.*

**Figure 2:**
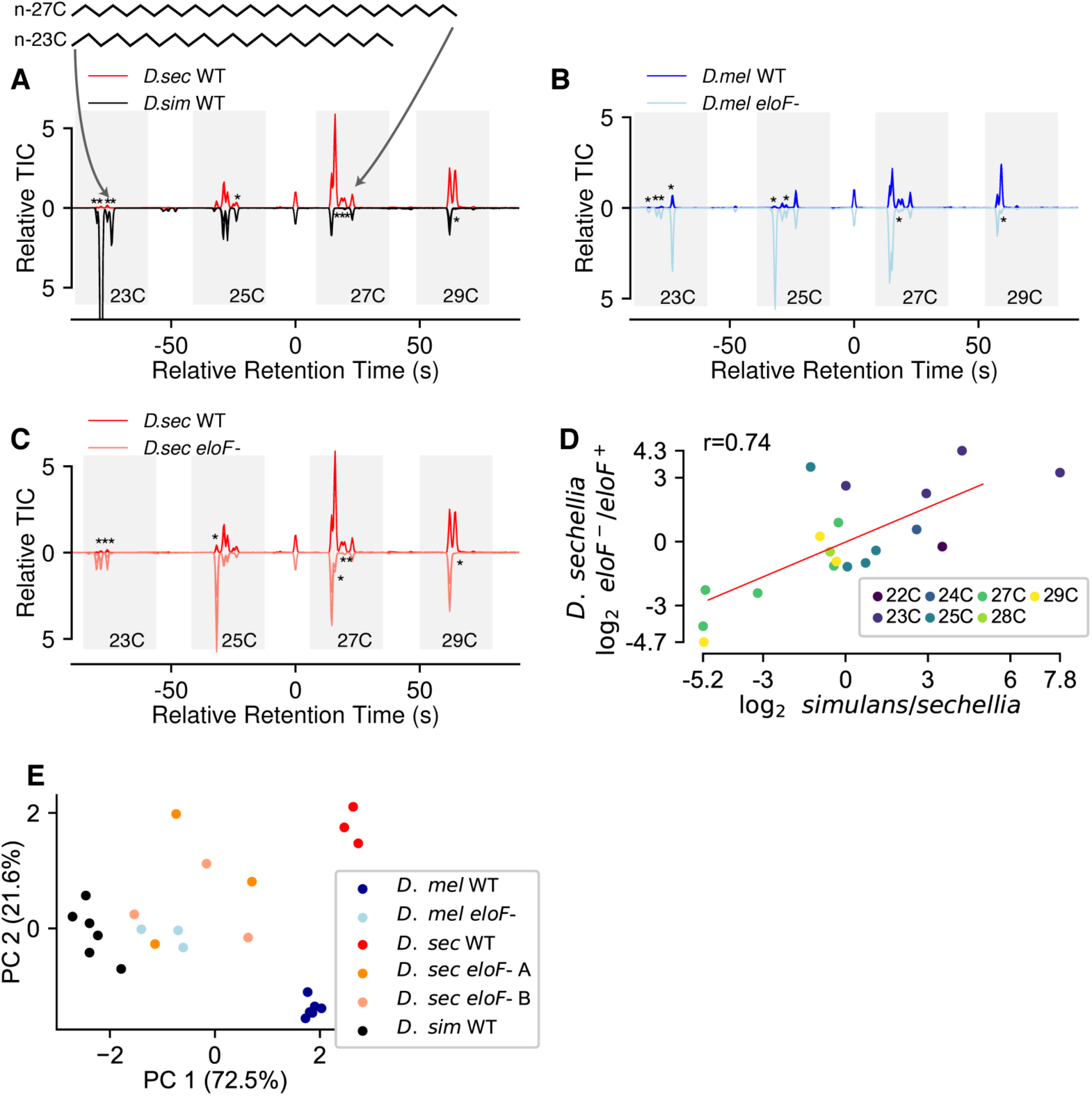
*eloF-* flies have an overall shorter CHC complement A) Total ion chromatographs of the hydrocarbon profile of wild-type *D. sechellia* (top) and *D. simulans* (bottom). Retention time and abundance is relative to the n-hexacosane (26C) normalization peak. Grey regions indicate number of carbons in CHC backbone. CHCs with more than a 3-fold change marked with asterisks at the location of the peak in the genotype with lower production. B-C) Total ion chromatographs of the hydrocarbon profile of wild-type (top) and *eloF-* (bottom) *D. melanogaster* (A) and *D. sechellia* (B). D) Average log2 fold changes of the measured compounds between *D. simulans* and *D. sechellia* vs log2 fold changes between wild-type and knockout of *eloF* in *D. sechellia*. Points are colored by the number of carbons in the backbone. E) Principal components analysis of wild-type and *eloF- D. melanogaster, simulans, and sechellia*. Principal components were calculated for the wild-type data, then *eloF-* data projected onto the same coordinates.

To explore the effects of our candidate genes on CHC profiles, we studied the phenotypic effects of their RNAi knockdowns in *D. melanogaster.* We did not pursue *desatF*, which already has a well-established role in *Drosophila* speciation (Legendre *et al.* 2008; Fang *et al.* 2009; Shirangi *et al.* 2009), or *FASN3*, which is essential for viability (Chung and Carroll 2015). For the remaining three CHC-related candidates, we created RNAi knockdowns in *D. melanogaster* females for each of these genes specifically in oenocytes by crossing PromE(800)-gal4 males with UAS-shRNA females from the TRiP project (Billeter *et al.* 2009; Perkins *et al.* 2015), then screened the CHC profiles of the progeny by GCMS. As negative controls, we crossed PromE(800)-gal4 males with females of Bloomington stock #32186, which carries 10 copies of UAS-driven mCD8-tagged GFP.

Of our three candidate genes, we found that one (*CG8534*) was essential for viability. Its highest expression is in the 3^rd^-4^th^ day of pupation (Graveley *et al.* 2011), so it may be involved in development. Attempts to delay induction of *gal4-*driven RNAi by incubating larvae at 18°C were not successful in rescuing females.

Knockdown of our second candidate (*bond*) in females led to ~60% increases in levels of pentacosadiene (a 25 carbon hydrocarbon) and ~60% decrease in levels of heptacosadiene (27 carbon) (Supplemental Figure 3). However other hydrocarbons were not significantly affected.

We observed the most pronounced effects for RNAi knockdown of our third candidate, *eloF*. We found that female flies with *eloF* knocked down have significantly fewer long-chain CHCs and more short-chain CHCs than wildtype flies (>3-fold change between CHCs with longer vs. shorter than 26 carbons; Figure 2B), consistent with previous work (Chertemps *et al.* 2007). Interestingly, *eloF* also had the strongest ASE among the six candidate genes (79-fold higher expression from *D. sechellia* alleles).

To examine the effect of *eloF* on CHCs in *D. sechellia*, we used CRISPR/Cas9 genome editing to create two independent lines of *D. sechellia* with *eloF* knocked out and replaced with P3-*RFP.* As expected, nearly all of the CHCs whose levels changed after *eloF* knockdown in *D. melanogaster* show a similar difference in *D. sechellia* (Figure 2C). Thus, we conclude that the molecular substrates and products of *eloF* are substantially similar between *D. melanogaster* and *sechellia.*

We noticed that there was a strong correlation between the changes observed between the sister species *D. simulans* and *D. sechellia* and the changes between wild-type and *eloF* depleted females from both *D. melanogaster* and *D. sechellia* (Figure 2D and Supplemental Figure 4). Consistent with *eloF*’s role as a fatty acid elongase, much of this variation consisted in broad differences in overall length of the hydrocarbons. To visualize entire CHC profiles, we performed principal components analysis, which showed that 94% of the total variation was captured by the first two components. The first principal component of variation separated *D. simulans* from both *D. melanogaster* and *D. sechellia* (Figure 2E). While knockdown or knockout of *eloF* did not completely transform the profiles of either species to *D. simulans*, it did make the profiles significantly closer. Thus, we concluded that one or more of the products of *eloF* may be acting as an anti-aphrodisiac to *D. simulans* males (or, alternatively, one of the substrates may be an aphrodisiac).

Notably, several previous studies have mapped quantitative trait loci (QTLs) that include *eloF*. For example, *eloF* is located within QTLs affecting CHC differences and mate discrimination between *D. simulans* and *D. sechellia* (Gleason *et al.* 2005; 2009), as well as a QTL for copulation frequency between *D. simulans* males and *D. mauritiana* females (Moehring *et al.* 2004). However in all of these studies, the QTLs also contained hundreds of other genes (including many other elongases). Therefore, although *eloF* is an excellent candidate gene, its role in reproductive isolation has not been explored.

### Expression of eloF is sufficient for species discrimination by *D. simulans* males

To determine whether the change in *eloF* expression (and concomitant CHC changes) could be responsible for sexual isolation between the species, we performed mate choice assays. We placed single *D. simulans* males in a chamber with a single female and recorded video in well-lit conditions for 30 minutes. We noted the time of the first instance of various copulatory behaviors, including tapping, male wing song, and licking (Figure 3A-C). With the exception of licking, these behaviors are not subject to rejection by females (the mating chambers are small enough that females are effectively unable to escape, while tapping is very rapid and wing song does not involve contact), and thus primarily represent choice by the males.

**Figure 3:**
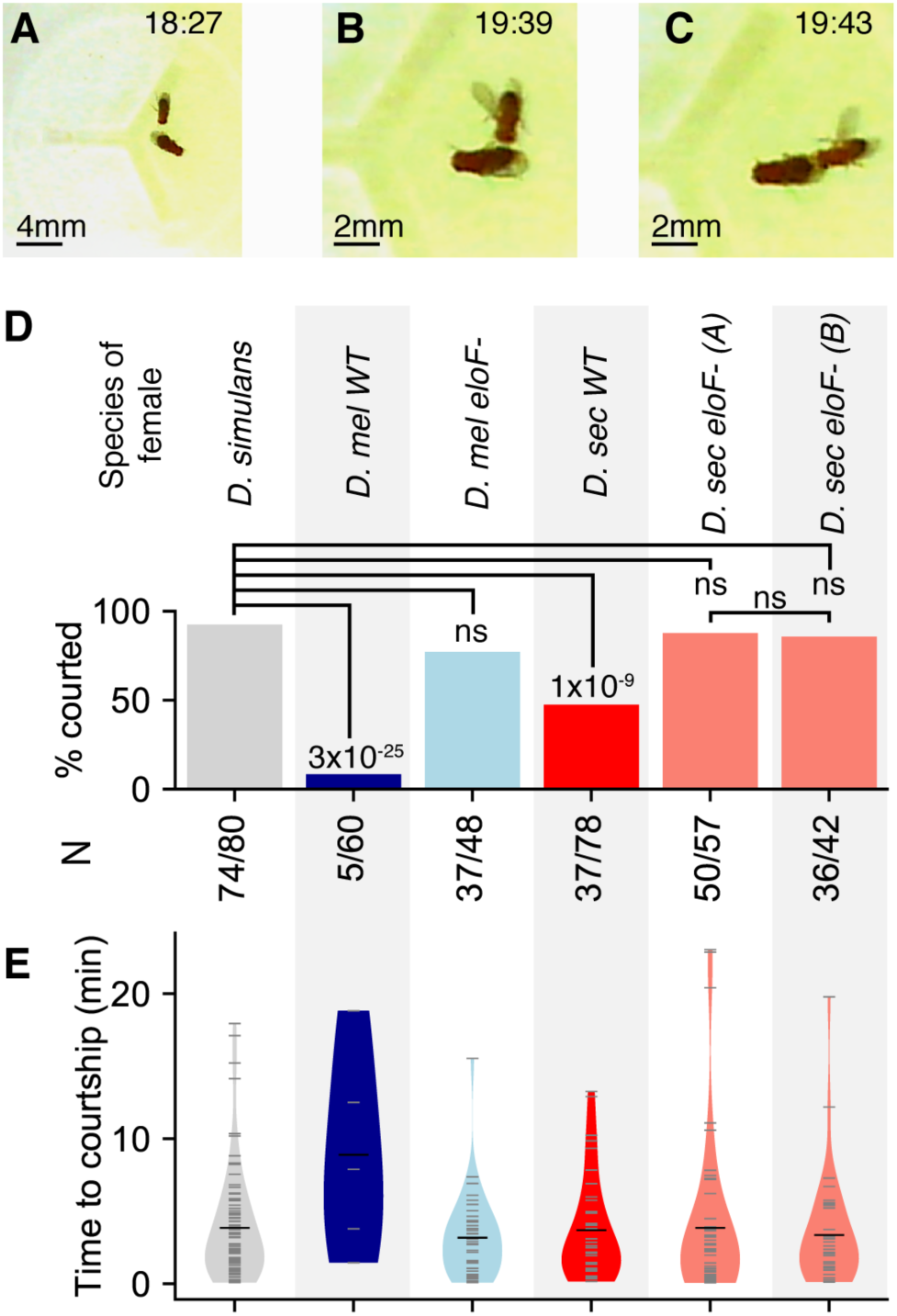
*D. simulans* males court interspecific *eloF*- females at significantly higher rates A-C) We recorded between 42 and 80 pairs of single *D. simulans* males courting single females of the indicated genotype. We recorded the time between male’s first tapping the female (and ostensibly sampling the female CHCs) and either singing behavior or licking of the female’s posterior prior to copulation. D) Female flies bearing a functional copy of *eloF* (*D. melanogaster* WT and *D. simulans* WT) were courted by *D. simulans* males at significantly lower rates than *D. simulans* conspecific females and interspecific females without *eloF*. We performed the indicated Fisher’s exact tests for differences in courtship rate (as measured by rate of proceeding to precopulatory licking), with Bonferroni-corrected *p*-values above each bar when significant. E) Violin plots of the delay between first contact between males and females and initiation of courtship. Black lines indicate mean time to courtship. Gray ticks indicate the underlying data. Although the *D. simulans* males were slower to court *D. melanogaster* WT females, this represents only 5 cases of courtship (out of 60 trials), and no comparisons were significant by t-test at even a nominal p=0.05 cutoff.

We first tested whether *eloF* might drive the behavioral isolation of *D. simulans* and *D. sechellia,* and so tested *D. sechellia* females with *D. simulans* males. As expected, *D. simulans* males courted wild-type *D. sechellia* females at a significantly lower rate than *D. simulans* females. Remarkably, *D. simulans* males courted *eloF- D. sechellia* females at the same rate as conspecific females (Figure 3D). We observed no significant difference in the courtship rate between the two independently generated *D. sechellia* knockout lines.

We then asked whether *eloF* might also mediate mate discrimination between *D. melanogaster* and *D. simulans.* As expected, when *D. simulans* males were presented with wildtype *D. melanogaster* females they rarely proceeded to courtship (Figure 3D and Supplemental Figure 5A). However, when we knocked down *eloF* expression in *D. melanogaster* females using oenocyte-specific RNAi, males courted them at rates only slightly lower than conspecifics.

The choice by males seems to be nearly binary. In the rare cases when *D. simulans* males did court wild-type *D. melanogaster* females, they did so approximately as quickly as they did for *D. simulans* females (Figure 3E and Supplemental Figure 5B). In none of the comparisons was there a significant difference in time between first contact between the flies and any of the steps in courtship at a nominal (i.e. without correcting for multiple testing) α=0.01 level.

## Discussion

Sexual selection in *Drosophila* has been studied for over one hundred years, with chemical odorants quickly being noticed as a primary signal (Sturtevant 1915), although the study of the evolution of these odorants came only after gas chromatography allowed the separation of different components (Hedin *et al.* 1972). Early work in the field sought to identify differences in CHC profiles between species and their effects on mating (Pechine *et al.* 1985; Jallon and David 1987; Cobb and Jallon 1990), and more recent genetic approaches have allowed for mapping of QTLs affecting these CHC differences (Moehring *et al.* 2004; Gleason *et al.* 2005; 2009). However, pinpointing the genes responsible for these changes is still quite difficult (Shirangi *et al.* 2009).

In this study, we have found that RNA-seq in F1 hybrids is a rapid, efficient means of identifying genes potentially involved in phenotypic divergence. Neither comparisons of expression across tissues nor of ASE within a single tissue was able to sufficiently narrow the list of candidate genes (Figure 1C); however, the combination of these orthogonal filters, together with gene annotations, allowed us to focus on only three excellent candidate genes. This can be compared with the most widely used alternative for studying the genetic basis of phenotypic divergence, QTL mapping. In QTL mapping, hundreds of progeny from genetic crosses must be genotyped and phenotyped, requiring years of effort even for rapidly reproducing species such as Drosophilids. Moreover, this effort leads to QTLs that typically span over a hundred genes, since resolution is limited by infrequent recombinations. Therefore, follow-up studies to test specific genes are often prohibitive. We envision that our approach of intersecting filters based only on RNA-seq in F1s may be widely applicable to other tissue-specific, sex-specific, stage-specific, or condition-specific traits that differ between interfertile populations or species.

Consistent with other recent observations (Shahandeh *et al.* 2018), we found that CHC differences between the species seem to be the major source of sexual isolation between *D. simulans* males and females from both *D. sechellia* and *D. melanogaster*, and we also showed that ablating *eloF* alleviates nearly all of the isolation from both *D. sechellia* and *D. melanogaster*. The magnitude of this effect is comparable to the reduction in barriers between *D. simulans* males and *D. melanogaster* females by ablating oenocytes entirely, a much more radical intervention (*eloF* appears to represent ~85% of the barrier in this study, compared to ~100% in Billeter *et al.* 2009).

One important caveat is that this isolation is observed under forced-choice laboratory conditions. Providing the choice between conspecifics and heterospecifics has been shown to increase isolation, while rates of hybridization in the wild have been strikingly higher than laboratory predictions (Coyne *et al.* 2005; Llopart *et al.* 2005).

Our identification of *eloF* as the necessary for *D. simulans* isolation is buttressed by understanding its role in the biochemical pathways of CHC synthesis but does not entirely depend on that foreknowledge. It is important that we were able to design our experiments knowing that the CHC biochemical pathway takes place almost completely in the oenocytes (Wicker-Thomas *et al.* 2015). However, having identified the candidate genes using RNA-seq, previous work investigating CHC synthesis allowed us to hypothesize why the candidates lead to different CHC profiles (Coyne 1996; Ferveur *et al.* 1997; Coyne *et al.* 1999; Labeur *et al.* 2002; Chertemps *et al.* 2007; Legendre *et al.* 2008). An interesting direction for future work would be to measure the effects of knocking out other genes in this pathway on CHC profiles and reproductive isolation.

Because *eloF* affects so many CHCs, it is not clear which CHC(s) act as the discriminative signal. The 27-carbon CHC 7,11-heptacosadiene has been shown to be involved in male *D. melanogaster* and *D. simulans* preference (Antony *et al.* 1985; Billeter *et al.* 2009), although other CHCs could also contribute. Further, the identity of the male receptor is unknown, although Gr32a seems to be the major chemoreceptor in *D. melanogaster* responsible for species recognition (Fan *et al.* 2013). While reagents in non-*melanogaster* Drosophilids are now available (Stern *et al,* 2017), screening multiple gustatory receptors in *D. simulans* is not yet as straightforward as an RNAi experiment in *D. melanogaster*.

However, even without knowing the specific causal CHCs we can hypothesize a parsimonious evolutionary scenario to explain our observations. *D. sechellia* and *D. melanogaster* both express *eloF* in female oenocytes; therefore this is likely to be the ancestral state for these species, with the 79-fold lower *eloF* expression in *D. simulans* being a derived change specific to this species. Our experiments show that *D. simulans* males prefer mates lacking *eloF*, suggesting that male preferences have co-evolved with CHC profiles in *D. simulans*. An intriguing question for future work will be whether the gene(s) responsible for this co-evolved male preference could be identified with a similar tissue-specific ASE approach as demonstrated here.

Another open question regards the sequence changes that have led to the expression differences of *eloF*. It seems significant that both a nearby coding gene (*CG8534*, also a fatty acid elongase) and a non-coding RNA (*CR44035*, of unknown function) share a similar pattern of female oenocyte-specific ASE. Neither of the genes bordering these 3 genes share this pattern, suggesting the existence of a species-variable topologically associated domain that is transcriptionally active in *D. sechellia* but not *D. simulans*. The transcription factor Doublesex has been implicated in the evolution of other *Drosophila* species’ CHC profiles (Shirangi *et al.* 2009), but searches for clear changes in canonical or non-canonical Doublesex binding sites have been fruitless in the species pair in this work. Further, the set of fixed changes is too large to easily test just a small set of candidates—in the noncoding region around *eloF* and *CG8534*, there are 136 SNPs and 10 indels (comprising 67 bases) where *D. simulans* has a derived allele differing from both *D. sechellia* and *D. melanogaster* (thus matching the parsimonious evolutionary scenario described above), in addition to several nonsynonymous changes in *eloF* (Supplemental Figure 6). An association study of *eloF* expression or CHC profiles in a panel of sequenced *D. simulans* may provide more targeted hypotheses, but only if the causal variant(s) are segregating within *D. simulans*, which seems unlikely given the major effect they would have on CHCs that are essential for mate choice.

Unlike previous observations that CHC changes can affect desiccation resistance (Chung *et al.* 2014; Ferveur *et al.* 2018), our preliminary tests of *eloF’s* effects on desiccation did not yield a strong effect (data not shown). These studies examined flies from widely varying ecological niches (Australian desert/jungle and France/Zimbabwe), whereas *D. simulans* and *D. sechellia* have overlapping ranges (Matute and Ayroles 2014). Thus, we would not expect strong pressure for differences in tolerance to desiccation.

Evolution of elongase expression may be involved in other insect speciation events as well. For instance, QTL studies between the jewel wasps *Nasonia vitripennis* and *N. giraulti* have implicated an elongase in CHC changes between those species (Niehuis *et al.* 2011). Furthermore, our analysis of CHC profiles in stingless bees shows at least two speciation events that show broad changes in the length of CHC backbones, which may be explained by divergence in elongase activity (Supplemental Figure 7; Nunes *et al,* 2017). Therefore, we hypothesize elongases may represent a general mechanism contributing to many cases of reproductive isolation in diverse insects.

## Materials and Methods

### RNA extraction and sequencing

Oenocyte and fat body dissections were performed as described in Krupp and Levine (2010). The oenocytes and fat body of 10-day-old *D. simulans/D. sechellia* hybrid flies were isolated separately from the dorsal abdominal segments of both adult male and female abdomens. Each tissue sample represented the pooled material collected from 20 flies. Hybrid flies were reared in a 12hr light:12 hr dark cycle and tissues dissected at equal time intervals across a 24hr period. Immediately following dissection tissues were placed into cell lysis buffer to aid in preserving the integrity of the RNA. Total RNA was isolated using the RNeasy Micro kit (Qiagen).

We prepared libraries from the RNA using the NextFLEX RNA-seq library preparation kit (BioO Scientific, Austin, TX), and sequenced the libraries using 101bp paired end reads on an Illumina HiSeq 2000.

We created a corrected *D. simulans* genome by using bowtie2 version 2.2.5 with arguments `--very-sensitive` to map genomic DNA reads from *D. simulans* and *D. sechellia* to the FlyBase 2.01 *D. simulans* reference genome (Hu *et al.* 2013; Coolon *et al.* 2014). Polymorphisms were called using GATK (`HaplotypeCaller --genotyping_mode DISCOVERY -fixMisencodedQuals -stand_emit_conf 10 -stand_call_conf 30`) (DePristo *et al.* 2011), then the ~34,000 SNPs that were fixed in both *D. simulans* and *D. sechellia* were replaced with the consensus sequence (this step was more important for creating a *simulans/sechellia* version of the *D. melanogaster* genome for Supplemental Figure N). RNA-seq reads were mapped to the reference genome using STAR with arguments `--outFilterMultimapNmax 1 --outSAMattributes MD NH --clip5pNbases 6 --sjdbGTFfile` (Dobin *et al.* 2013). Following the WASP pipeline, duplicate reads were discarded randomly, then filtered based on whether reads with the alleles swapped *in silico* to create artificial transcripts from the other species mapped to the same position (van de Geijn *et al.* 2015). Reads were assigned to a species only if both paired ends mapped unambiguously to one species, and allele-specific expression negative binomial p-values were calculated from aligned read counts using DESeq2 with model ~Replicate + AlignsToSpecies (Love *et al.* 2014). Default DESeq settings were used to correct for multiple hypothesis testing. Transcript abundances were estimated using kallisto with default arguments (Bray *et al.* 2016). We used sleuth to identify differentially expressed genes between samples with matched sex and tissue type (Pimentel *et al.* 2017).

### Fly rearing

For RNAi flies, virgin females of the shRNA driver were isolated within 18 hours of eclosion, then kept isolated from males for 3 days on standard cornmeal media to ensure virgin status. We used Bloomington Stock IDs 34676 (*bond)*, 53947 (*eloF),* 53299 (*CG8534),* and 32186 (GFP control). We combined approximately 25 UAS-shRNA females with approximately 10 Gal4 driver males. Adults were moved to fresh vials every 3 days to ensure separation of the parents and the Gal4+UAS offspring.

Knockout *D. sechellia* flies were created using CRISPR/Cas9 mediated editing. We designed guides to cut at the 55^th^ nucleotide downstream of the ATG and the 114^th^ nucleotide upstream of the stop codon of *GM23846* (the *D. sechellia* ortholog of *eloF)*. We used sense oligos CTTCGCAGCGATCCATGGGTCCCCA (gene 5’-ward cut site) and CTTCGATCCGCATCCGTAGGTCAA (gene 3’-ward cut site). Embryos were injected (WellGenetics, Taipei, Taiwan) with both guides and a dsDNA donor containing ~1000bp homology arms and RFP driven by 3 P3 promoters and flanked by LoxP sites. Embryos were from the *D. sechellia* genome strain #14021-0248.25.

All flies, either RNAi or CRISPR edited were separated by sex within 18 hours of eclosion, then kept isolated for 5-7 days to ensure virgin status. Any vials with larvae after 5 days were discarded. Since the PromE(800)-gal4 construct is balanced with Tm3.5b, we selected straight-winged flies as RNAi positive.

### Gas chromatography–mass spectrometry

We performed GCMS by anesthetizing 5 females at 4°C for 3-5 minutes, then washing them for 5 minutes with 50µL of hexane spiked with 10mg/mL of n-hexane as a standard. Spectra were obtained using an Agilent (HP) 7890/5975 single quadrupole GC-MS instrument with a split ratio of 1:20, injector temperature of 280°C, and an oven temperature program of 35°C hold for 3.75min, 20°C/min ramp from 35°C to 320°C, and a 320°C hold for 7 min. We collected spectra for at least 3 sets of 5 flies for each genotype. Identities of different hydrocarbon peaks were inferred by inspecting the singly-ionized mass spectrum bin.

### Mating assays

We performed mating assays by anesthetizing separate vials of males and females at 4°C for 3-5 minutes, then used a paintbrush to transfer one male and one female to each well of the mating chamber. The mating chamber was 3D printed from acrylic plastic and has 18 separate 2cm diameter × 5mm circular wells, with a removable clear plastic lid. We allowed flies to acclimate at room temperature and ambient light for 10-15 minutes, then recorded 30m of video with bright lights, which we found were required for *D. simulans* males to initiate courtship. The mating light was a 75W, 14” circular fluorescent bulb placed approximately 30cm above the mating chamber. Video of mating assays was recorded using a Dino-Lite digital microscope, then analyzed by two separate graders (PAC and NMK), who recorded the time of first contact by the male, the time of the male first following the female, the time of the first wing song by the male, and the time of first licking by the male of the female’s abdomen (Sokolowski 2001). Graders were blinded to the fly identities in each video.

### Data Availability

Sequencing data has been deposited at the Gene Expression Omnibus under access number GSE114478. An interactive tool to explore the RNA-seq dataset is available at http://combsfraser-oenocytes.appspot.com/.

## Acknowledgements

We thank Nirao Shah and Osama Ahmed for helpful discussions on mating assays. We thank David Stern for helpful discussions and the *D. simulans* tsimbazaza strain. The Bloomington Drosophila Stock Center (NIH P40OD018537) provided the RNAi strains and the Drosophila Species Stock Center at UCSD (now at Cornell University) provided the D. sechellia strains through support of NSF CSBR grant 1351502. GCMS was performed at the Vincent Coates Foundation Mass Spectrometry Laboratory, Stanford University Mass Spectrometry (http://mass-spec.stanford.edu), which is supported in part by NIH P30 CA124435. This work was supported by NIH grant 2R01GM097171-05A1.

## Supplemental Information

**Supplemental Table 1:** RNA sequencing library details. Unique deduplicated mapped reads indicates reads that pass the WASP pipeline (van de Geijn *et al*. 2015), i.e. that map to a single position and map to the same position when alleles are swapped. Overall mapping rates (including multimappers) are typically around 65%.

| TISSUE | SEX | REPLICATE | TOTAL<br>PAIRED<br>END READS | UNIQUE DEDUPLICATED<br>DE-BIASED READS (%) | READS ASSIGNABLE<br>TO PARENT<br>(% OF MAPPED) | D. SECHELLIA<br>READS<br>(% OF ASSIGNABLE) |
| --- | --- | --- | --- | --- | --- | --- |
| Fat body | Female | 1 | 29,958,590 | 3,099,853 (10.3%) | 1,166,014 (37.6%) | 683,982 (58.7%) |
| Fat body | Female | 2 | 26,693,093 | 4,156,810 (15.6%) | 1,610,951 (38.8%) | 919,575 (57.1%) |
| Fat body | Male | 1 | 30,135,650 | 2,671,563 (8.9%) | 917,125 (34.3%) | 502,862 (54.8%) |
| Fat body | Male | 2 | 36,340,739 | 4,283,897 (11.8%) | 1,726,324 (40.3%) | 910,829 (52.8%) |
| Oenocyte | Female | 1 | 48,560,642 | 5,189,309 (10.7%) | 1,711,402 (33.0%) | 994,986 (58.1%) |
| Oenocyte | Female | 2 | 41,310,971 | 4,295,068 (10.4%) | 1,541,471 (35.9%) | 832,732 (54.0%) |
| Oenocyte | Male | 1 | 26,683,030 | 2,540,477 (9.5%) | 808,148 (31.8%) | 408,929 (50.6%) |
| Oenocyte | Male | 2 | 53,693,579 | 4,324,451 (8.1%) | 1,478,937 (34.2%) | 790,366 (53.4%) |

**Supplemental Figure 1:**
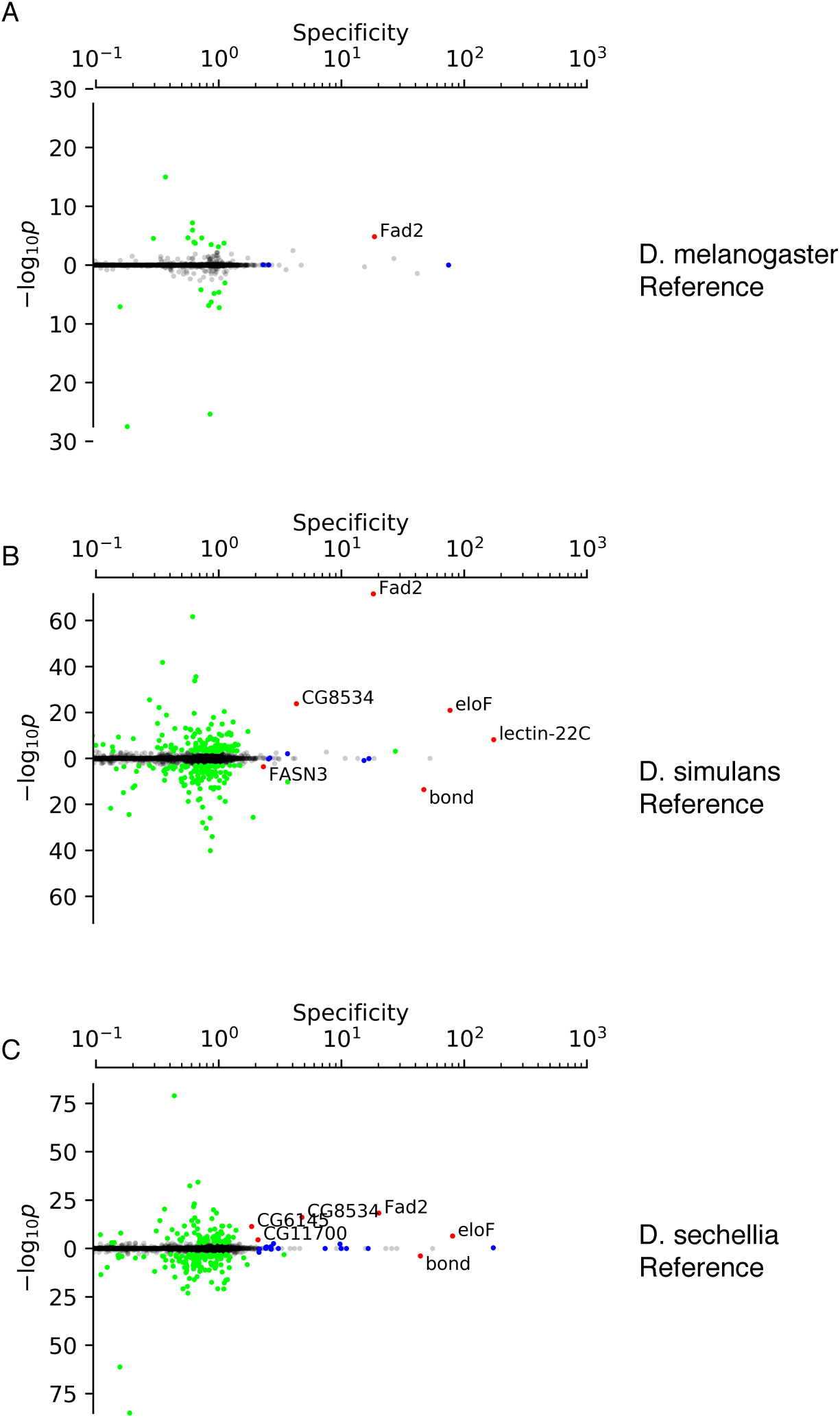
Primary candidate genes are robust to the choice of reference genome. Specificity-ASE plot as in Figure 1B, but with reads mapped to the (A) *D. melanogaster,* (B—same as Fig. 1B) *D. simulans,* or (C) *D. sechellia* genome.

**Supplemental Figure 2:**
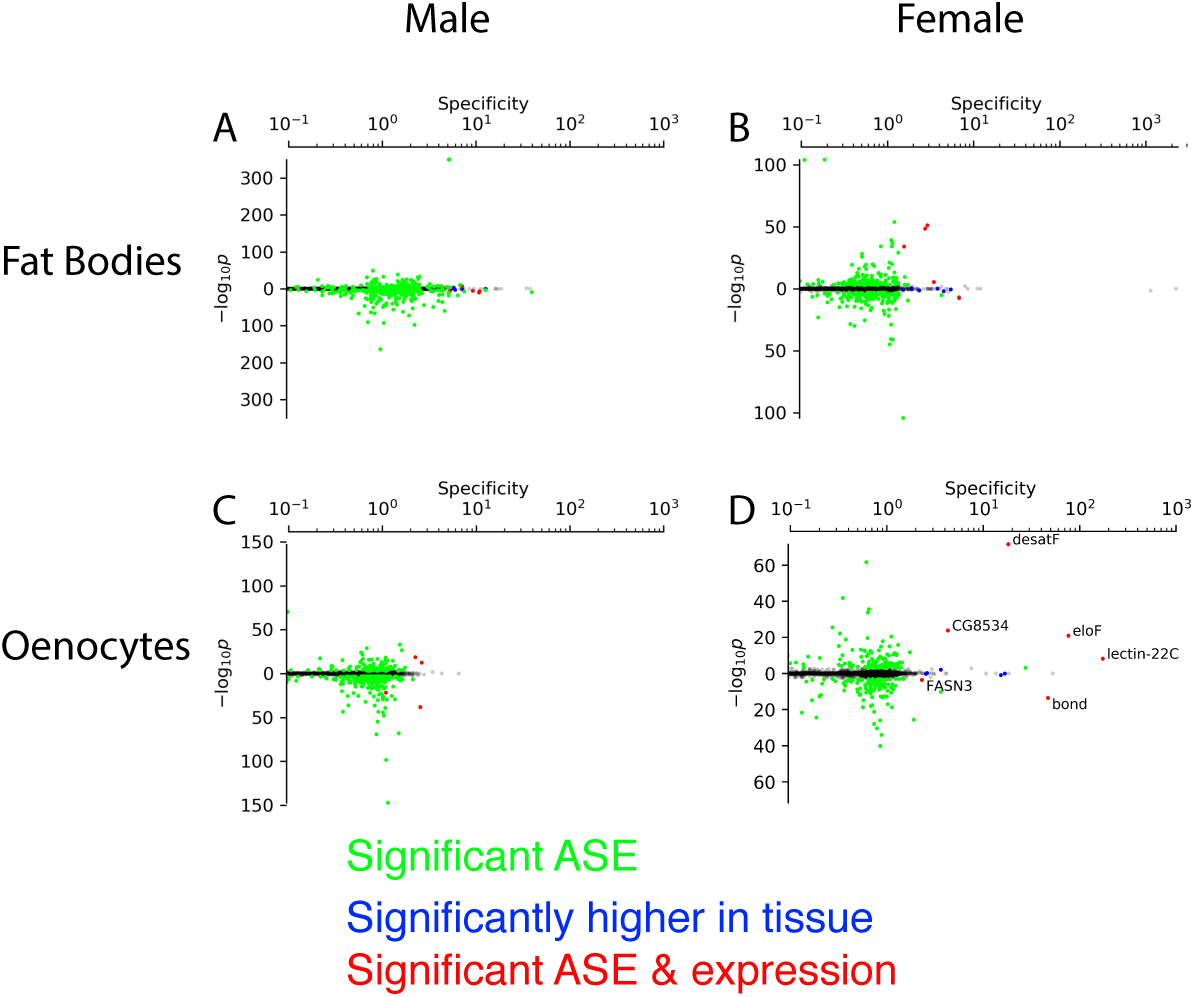
Male oenocytes do not show strong tissue-specific expression. **A-C)** ASE and tissue specificity as in Figure 1B, except for male fat bodies (A), female fat bodies (B), male oenocytes (C), and female oenocytes (D—same as Fig. 1B).

**Supplemental Table 2:** References for Gene Ontology Terms in Table 1 Summary of FlyBase-curated function gene ontology codes and evidence for genes with female oenocyte-spcific and allele-specific expression (Gramates *et al.* 2017). Direct evidence indicates evidence codes of “Inferred from Direct Assay”, “Inferred from Genetic Interaction”, and “Inferred from Physical Interaction”.

| *Gene* | *GO Term* | *GO Term ID* | *Direct evidence?* | *Reference* |
| --- | --- | --- | --- | --- |
| *eloF* | fatty acid elongase activity | GO:0009922 | Y | (Chertemps *et al.* 2007) |
| *desatF* | stearoyl-CoA 9-desaturase activity | GO:0004768 | N | (Finn *et al.* 2017; Gramates *et al.* 2017) |
| *CG8534* | fatty acid elongase activity | GO:0009922 | N | (Szafer-Glusman *et al.* 2008) |
| *FASN3* | 3-oxoacyl-[acyl-carrier-protein] synthase activity | GO:0004315 | N | (Finn *et al.* 2017; Gramates *et al.* 2017) |
| *lectin-22C* | Galactose binding; Carbohydrate binding | GO:0005534; GO:0030246 | N | (Theopold *et al.* 1999; Tanji *et al.* 2006) |
| *bond* | fatty acid elongase activity | GO:0009922 | Y | (Szafer-Glusman *et al.* 2008) |

**Supplemental Figure 3:**
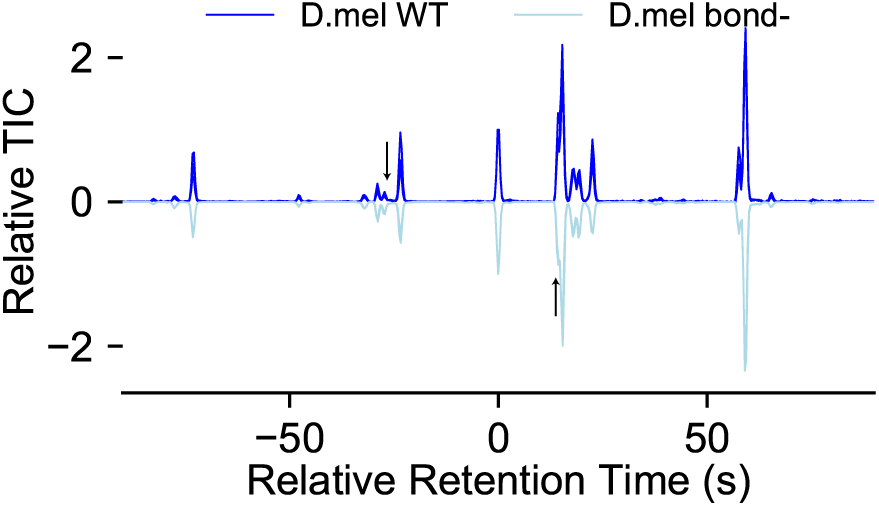
GCMS of *bond*- *D. melanogaster* females. Arrows indicate hydrocarbons with ~60% change in levels.

**Supplemental Figure 4:**
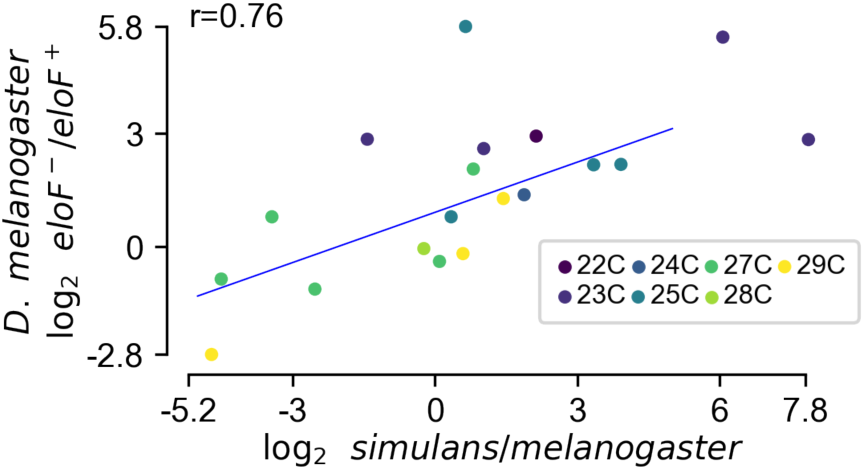
Pairwise comparison of *D. melanogaster* WT vs *eloF* knockdown and *D. melanogaster* WT vs *D. simulans* WT. Log fold change between the indicated comparisons for average area under the total ion chromatogram curve of each of 19 different CHC peaks.

**Supplemental Figure 5:**
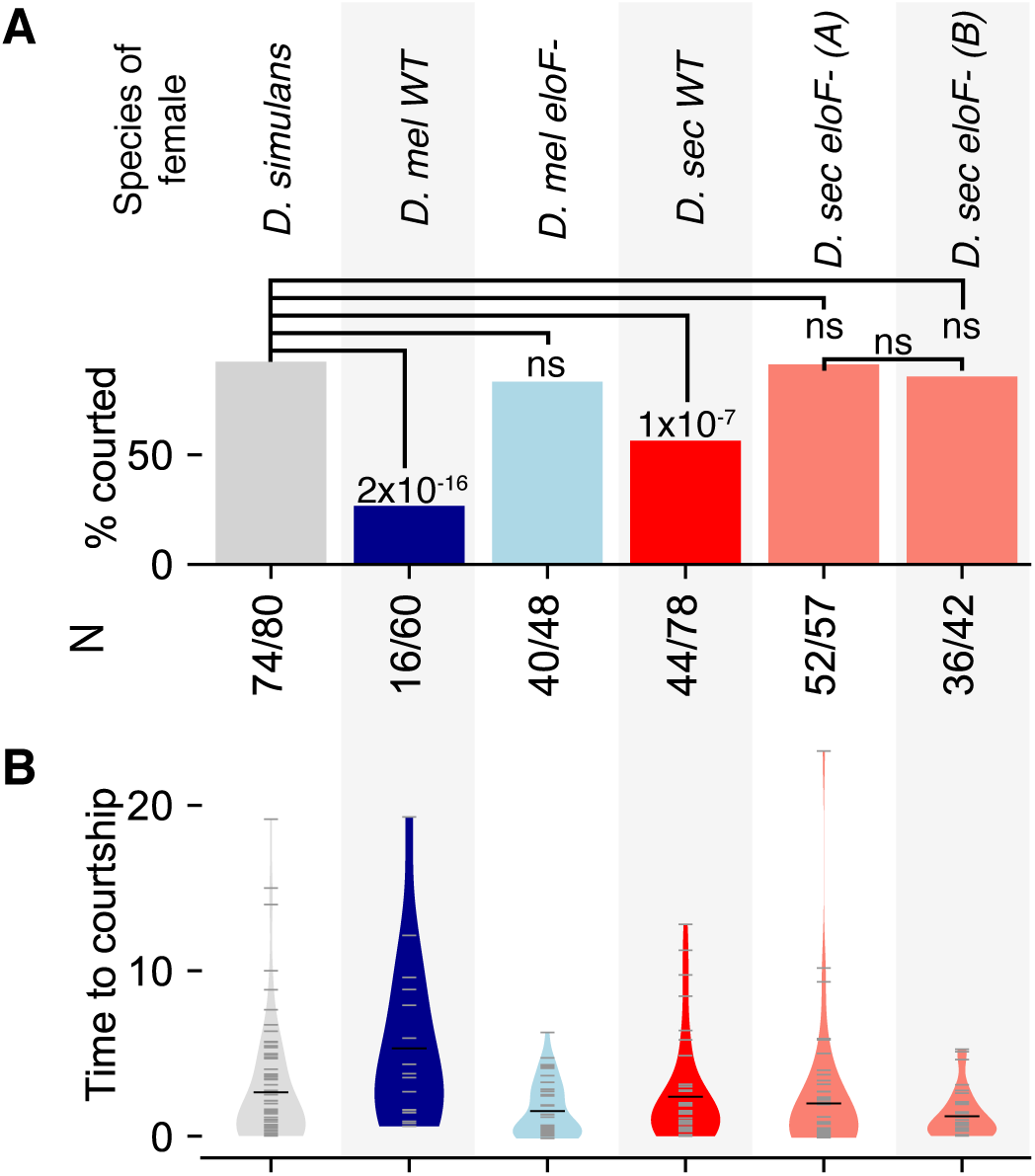
*D. simulans* males perform wing song for *eloF-* females at significantly higher rates than *eloF+* females. Courtship rate (A) and delay until initiation of wing song (B), as in Figure 3D-E, except using time to initiation of wing song instead of pre-copulatory licking.

**Supplemental Figure 6:**
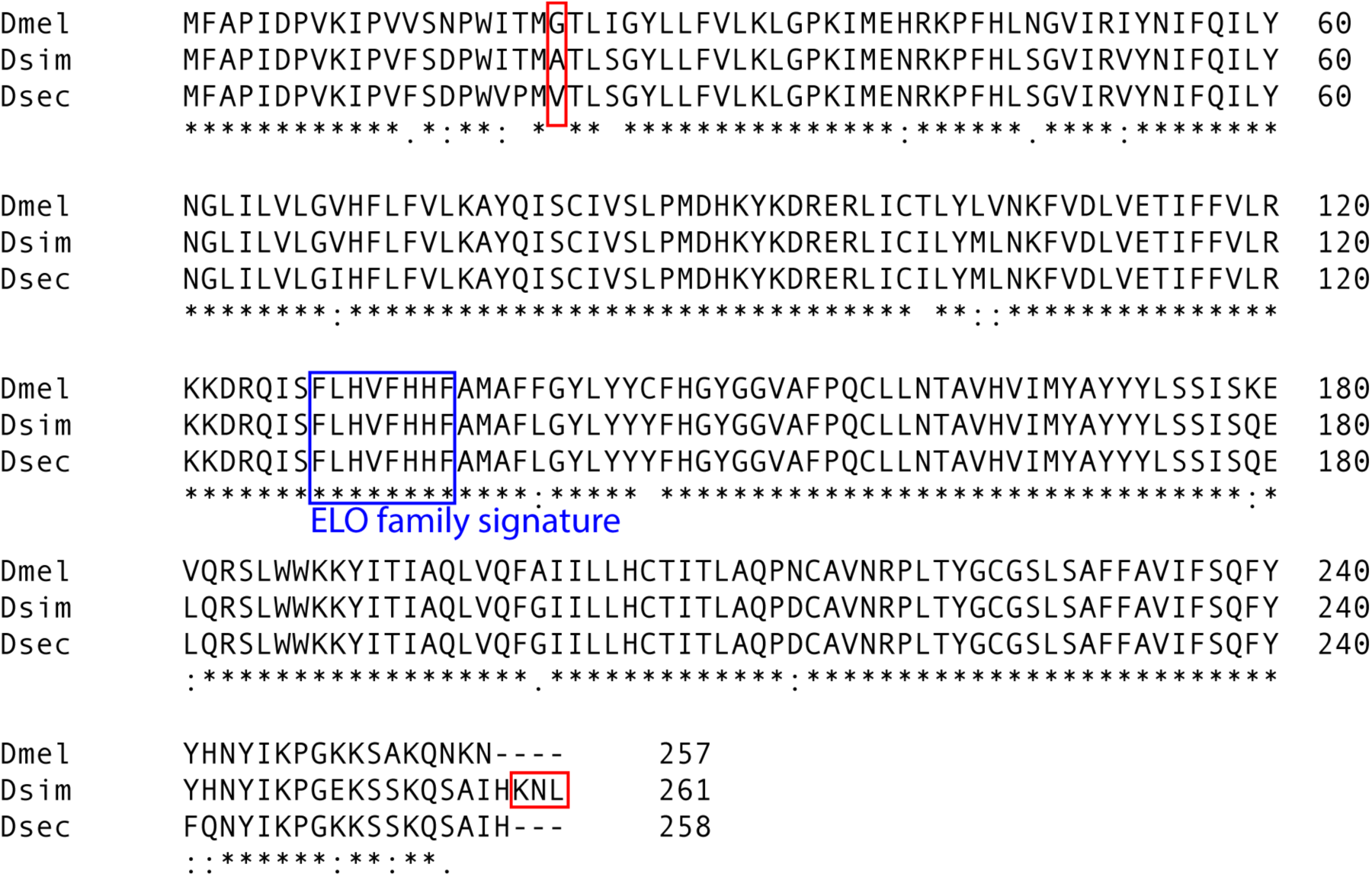
Protein alignments of the *D. melanogaster, simulans,* and *sechellia* versions of *eloF* show few *D. simulans-*specific changes. We aligned the coding sequences of eloF and its orthologs in *D. simulans* and *D. sechellia* using Clustal Omega (Sievers, *et al. 2012*). Red boxes indicate where the *D. simulans* does not match at least one of the other species. The blue box is the ELO family signature PS01188 from ProSite (Sigrist, *et al.* 2012).

**Supplemental Figure 7:**
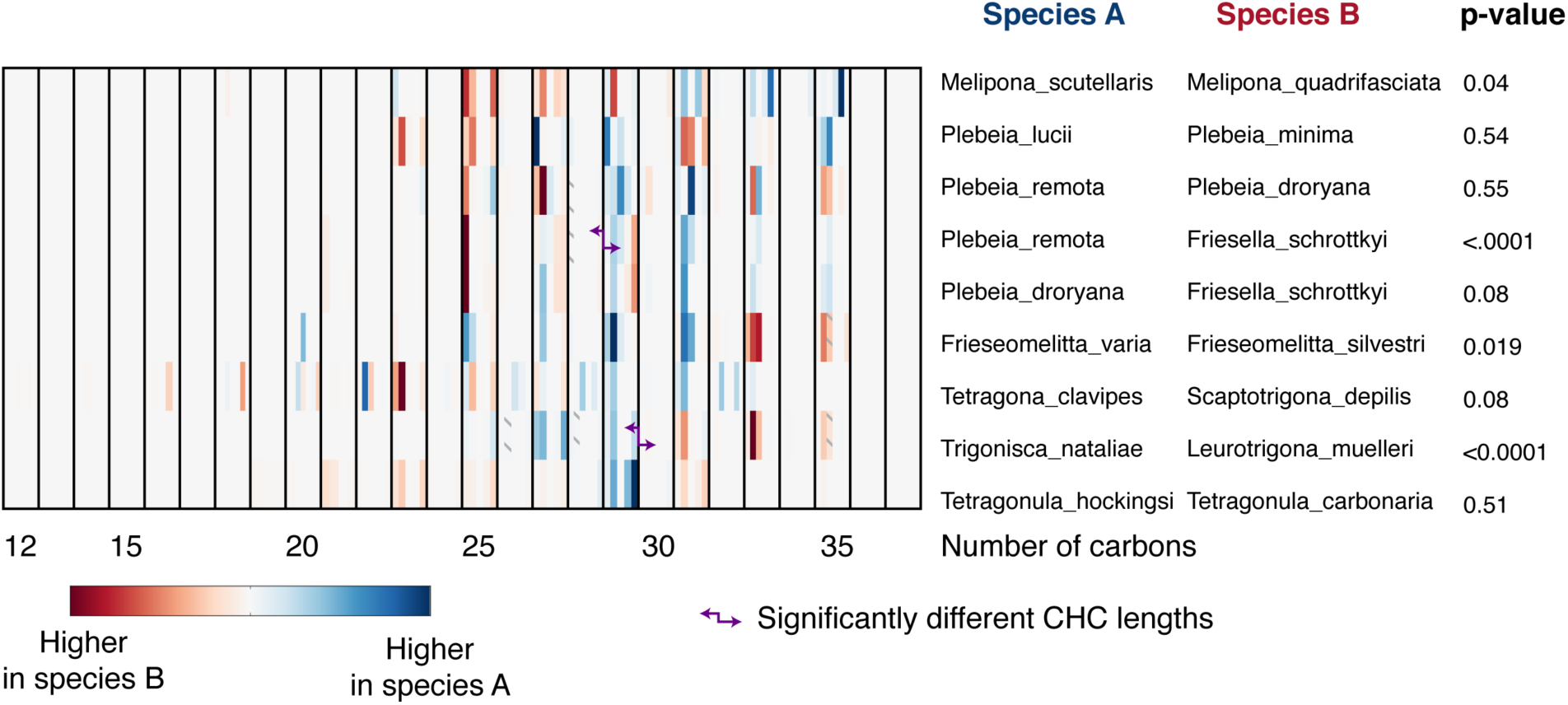
Hydrocarbon length is connected to speciation in some stingless bees. We examined the changes in hydrocarbon profiles of stingless bee queens between recently diverged species pairs, as measured in Nunes *et al*, 2017. To determine whether there was a change in overall hydrocarbon length, we looked for a critical CHC length that maximized the sum of the squares of CHCs shorter than the critical length plus the sum of the squares of CHCs longer than the critical length. To calculate p-values, we randomized the order of CHCs (while keeping CHCs with the same number of carbons together), performing 10,000 permutations. After Bonferroni correction, we found a significant divergence in CHC length between *P. remota* and *F. schrottkyi*, with *P. remota* having more CHCs with 29 or more carbons, and between *T. nataliae* and *L. muelleri*, with *L. muelleri* having more CHCs with 30 or more carbons.

## References

1. Antony, C., T. L. Davis, D. A. Carlson, J. M. Pechine, and J.-M. Jallon, 1985 Compared behavioral responses of male *Drosophila melanogaster* (Canton S) to natural and synthetic aphrodisiacs. J. Chem. Ecol. 11: 1617–1629.

2. Billeter, J.-C., J. Atallah, J. J. Krupp, J. G. Millar, and J. D. Levine, 2009 Specialized cells tag sexual and species identity in Drosophila melanogaster. Nature 461: 987–991.

3. Boyle, E. I., S. Weng, J. Gollub, H. Jin, D. Botstein et al., 2004 GO::TermFinder--open source software for accessing Gene Ontology information and finding significantly enriched Gene Ontology terms associated with a list of genes. Bioinformatics 20: 3710–3715.

4. Bray, N. L., H. Pimentel, P. Melsted, and L. Pachter, 2016 Near-optimal probabilistic RNA-seq quantification. Nat Biotech 34: 525–527.

5. Byrne, P. G., and W. R. Rice, 2006 Evidence for adaptive male mate choice in the fruit fly Drosophila melanogaster. Proc. Biol. Sci. 273: 917–922.

6. Chertemps, T., L. Duportets, C. Labeur, R. Ueda, K. Takahashi et al., 2007 A female-biased expressed elongase involved in long-chain hydrocarbon biosynthesis and courtship behavior in Drosophila melanogaster. Proc. Natl. Acad. Sci. 104: 4273–4278.

7. Chung, H., and S. B. Carroll, 2015 Wax, sex and the origin of species: Dual roles of insect cuticular hydrocarbons in adaptation and mating. Bioessays 37: 822–830.

8. Chung, H., D. W. Loehlin, H. D. Dufour, K. Vaccarro, J. G. Millar et al., 2014 A single gene affects both ecological divergence and mate choice in Drosophila. Science 343: 1148–1151.

9. Cobb, M., and J.-M. Jallon, 1990 Pheromones, mate recognition and courtship stimulation in the Drosophila melanogaster species sub-group. Anim Behav 39: 1058–1067.

10. Coolon, J. D., C. J. McManus, K. R. Stevenson, B. R. Graveley, and P. J. Wittkopp, 2014 Tempo and mode of regulatory evolution in Drosophila. Genome Res 24: 797–808.

11. Coyne, J. A., 1996 Genetics of differences in pheromonal hydrocarbons between Drosophila melanogaster and D. simulans. Genetics 143: 353–364.

12. Coyne, J. A., and H. A. Orr, 1997 “Patterns of Speciation in *Drosophila*” Revisited. Evolution 51:295–303.

13. Coyne, J. A., and H. A. Orr, 2004 Speciation. Sinauer Associates Incorporated.

14. Coyne, J. A., A. P. Crittenden, and K. Mah, 1994 Genetics of a pheromonal difference contributing to reproductive isolation in Drosophila. Science 265: 1461–1464.

15. Coyne, J. A., S. Elwyn, and E. Rolán-Alvarez, 2005 Impact of experimental design on Drosophila sexual isolation studies: direct effects and comparison to field hybridization data. Evolution 59: 2588–2601.

16. Coyne, J. A., C. Wicker-Thomas, and J.-M. Jallon, 1999 A gene responsible for a cuticular hydrocarbon polymorphism in Drosophila melanogaster. Genet. Res. 73: 189–203.

17. DePristo, M. A., E. Banks, R. Poplin, K. V. Garimella, J. R. Maguire et al., 2011 A framework for variation discovery and genotyping using next-generation DNA sequencing data. Nat Genet 43:491–498.

18. Dobin, A., C. A. Davis, F. Schlesinger, J. Drenkow, C. Zaleski et al., 2013 STAR: ultrafast universal RNA-seq aligner. Bioinformatics 29: 15–21.

19. Edward, D. A., and T. Chapman, 2011 The evolution and significance of male mate choice. Trends Ecol. Evol. (Amst.) 26: 647–654.

20. Fan, P., D. S. Manoli, O. M. Ahmed, Y. Chen, N. Agarwal et al., 2013 Genetic and neural mechanisms that inhibit Drosophila from mating with other species. Cell 154: 89–102.

21. Fang, S., C.-T. Ting, C.-R. Lee, K.-H. Chu, C.-C. Wang et al., 2009 Molecular evolution and functional diversification of fatty acid desaturases after recurrent gene duplication in Drosophila. Molecular Biology and Evolution 26: 1447–1456.

22. Ferveur, J.-F., J. Cortot, K. Rihani, M. Cobb, and C. Everaerts, 2018 Desiccation resistance: effect of cuticular hydrocarbons and water content in Drosophila melanogaster adults. PeerJ 6: e4318.

23. Ferveur, J. F., F. Savarit, C. J. O’Kane, G. Sureau, R. J. Greenspan et al., 1997 Genetic feminization of pheromones and its behavioral consequences in Drosophila males. Science 276:1555–1558.

24. Finn, R. D., T. K. Attwood, P. C. Babbitt, A. Bateman, P. Bork et al., 2017 InterPro in 2017-beyond protein family and domain annotations. Nucleic Acids Research 45: D190–D199.

25. Fowler, K., and L. Partridge, 1989 A cost of mating in female fruitflies. Nature 338: 760–761.

26. Garrigan, D., S. B. Kingan, A. J. Geneva, P. Andolfatto, A. G. Clark et al., 2012 Genome sequencing reveals complex speciation in the Drosophila simulans clade. Genome Res 22: 1499–1511.

27. Gleason, J. M., J.-M. Jallon, J.-D. Rouault, and M. G. Ritchie, 2005 Quantitative trait loci for cuticular hydrocarbons associated with sexual isolation between Drosophila simulans and D. sechellia. Genetics 171: 1789–1798.

28. Gleason, J. M., R. A. James, C. Wicker-Thomas, and M. G. Ritchie, 2009 Identification of quantitative trait loci function through analysis of multiple cuticular hydrocarbons differing between Drosophila simulans and Drosophila sechellia females. Heredity (Edinb) 103: 416–424.

29. Gramates, L. S., S. J. Marygold, G. D. Santos, J.-M. Urbano, G. Antonazzo et al., 2017 FlyBase at 25: looking to the future. Nucleic Acids Research 45: D663–D671.

30. Graveley, B. R., A. N. Brooks, J. W. Carlson, M. O. Duff, J. M. Landolin et al., 2011 The developmental transcriptome of Drosophila melanogaster. Nature 471: 473–479.

31. Greenspan, R. J., and J. F. Ferveur, 2000 Courtship in Drosophila. Annu. Rev. Genet. 34: 205–232.

32. Hedin, P. A., C. S. Niemeyer, R. C. Gueldner, and A. C. Thompson, 1972 A gas chromatographic survey of the volatile fractions of twenty species of insects from eight orders. J. Insect Physiol.18: 555–564.

33. Hu, T. T., M. B. Eisen, K. R. Thornton, and P. Andolfatto, 2013 A second-generation assembly of the Drosophila simulans genome provides new insights into patterns of lineage-specific divergence. Genome Res 23: 89–98.

34. Jallon, J.-M., and J. R. David, 1987 Variation in Cuticular Hydrocarbons Among the Eight Species of the *Drosophila melanogaster* Subgroup. Evolution 41: 294–302.

35. Kliman, R. M., P. Andolfatto, J. A. Coyne, F. Depaulis, M. Kreitman et al., 2000 The Population Genetics of the Origin and Divergence of the Drosophila simulans Complex Species. Genetics 156:1913–1931.

36. Labeur, C., R. Dallerac, and C. Wicker-Thomas, 2002 Involvement of desat1 gene in the control of Drosophila melanogaster pheromone biosynthesis. Genetica 114: 269–274.

37. Lachaise, D., J. R. David, F. Lemeunier, and L. Tsacas, 1986 The reproductive relationships of Drosophila sechellia with D. mauritiana, D. simulans, and D. melanogaster from the Afrotropical region. Evolution.

38. Lasbleiz, C., J.-F. Ferveur, and C. Everaerts, 2006 Courtship behaviour of Drosophila melanogaster revisited. Anim Behav 72: 1001–1012.

39. Lazareva, A. A., G. Roman, W. Mattox, P. E. Hardin, and B. Dauwalder, 2007 A role for the adult fat body in Drosophila male courtship behavior. PLoS Genet 3: e16.

40. Legendre, A., X.-X. Miao, J.-L. Da Lage, and C. Wicker-Thomas, 2008 Evolution of a desaturase involved in female pheromonal cuticular hydrocarbon biosynthesis and courtship behavior in Drosophila. Insect Biochem. Mol. Biol. 38: 244–255.

41. Llopart, A., D. Lachaise, and J. A. Coyne, 2005 An anomalous hybrid zone in Drosophila. Evolution 59: 2602–2607.

42. Love, M. I., W. Huber, and S. Anders, 2014 Moderated estimation of fold change and dispersion for RNA-seq data with DESeq2. Genome Biol 15: 31–21.

43. Matute, D. R., and J. F. Ayroles, 2014 Hybridization occurs between Drosophila simulans and D. sechellia in the Seychelles archipelago. J. Evol. Biol. 27: 1057–1068.

44. Moehring, A. J., J. Li, M. D. Schug, S. G. Smith, M. deAngelis et al., 2004 Quantitative trait loci for sexual isolation between Drosophila simulans and D. mauritiana. Genetics 167: 1265–1274.

45. Niehuis, O., J. Büllesbach, A. K. Judson, T. Schmitt, and J. Gadau, 2011 Genetics of cuticular hydrocarbon differences between males of the parasitoid wasps Nasonia giraulti and Nasonia vitripennis. Heredity (Edinb) 107: 61–70.

46. Noor, M. A., 1995 Speciation driven by natural selection in Drosophila. Nature 375: 674–675.

47. Partridge, L., 1980 Mate choice increases a component of offspring fitness in fruit flies. ature, Lond. Nature 283: 290–291.

48. Partridge, L., and M. Farquhar, 1981 Sexual activity reduces lifespan of male fruitflies. Nature 294:580–582.

49. Pechine, J. M., F. Perez, C. Antony, and J.-M. Jallon, 1985 A further characterization of Drosophila cuticular monoenes using a mass spectrometry method to localize double bonds in complex mixtures. Anal. Biochem. 145: 177–182.

50. Perkins, L. A., L. Holderbaum, R. Tao, Y. Hu, R. Sopko et al., 2015 The Transgenic RNAi Project at Harvard Medical School: Resources and Validation. Genetics 201: 843–852.

51. Phadnis, N., E. P. Baker, J. C. Cooper, K. A. Frizzell, E. Hsieh et al., 2015 The Drosophila melanogaster hybrid male rescue gene causes inviability in male and female species hybrids. Science 350:1552–1555.

52. Pimentel, H., N. L. Bray, S. Puente, P. Melsted, and L. Pachter, 2017 Differential analysis of RNA-seq incorporating quantification uncertainty. Nat Meth 14: 687–690.

53. Pischedda, A., M. P. Shahandeh, W. G. Cochrane, V. A. Cochrane, and T. L. Turner, 2014 Natural variation in the strength and direction of male mating preferences for female pheromones in Drosophila melanogaster. PLoS ONE 9: e87509.

54. Quinn, T. P., M. J. Unwin, and M. T. Kinnison, 2000 Evolution of temporal isolation in the wild: genetic divergence in timing of migration and breeding by introduced chinook salmon populations. Evolution 54: 1372–1385.

55. Sawamura, K., M.-T. Yamamoto, and T. K. Watanabe, 1993 Hybrid lethal systems in the Drosophila melanogaster species complex. II. The Zygotic hybrid rescue (Zhr) gene of D. melanogaster. Genetics 133: 307–313.

56. Servedio, M. R., 2007 Male versus female mate choice: sexual selection and the evolution of species recognition via reinforcement. Evolution 61: 2772–2789.

57. Servedio, M. R., and M. A. F. Noor, 2003 The Role of Reinforcement in Speciation: Theory and Data. Annu. Rev. Ecol. Evol. Syst. 34: 339–364.

58. Shahandeh, M. P., A. Pischedda, and T. L. Turner, 2018 Male mate choice via cuticular hydrocarbon pheromones drives reproductive isolation between Drosophila species. Evolution 72: 123–135.

59. Shirangi, T. R., H. D. Dufour, T. M. Williams, and S. B. Carroll, 2009 Rapid evolution of sex pheromone-producing enzyme expression in Drosophila. PLoS Biol 7: e1000168.

60. Sigrist, C. J. A., E. de Castro, L. Cerutti, B. A. Cuche, N. Hulo et al., 2012 New and continuing developments at PROSITE. Nucleic Acids Research 41: D344–D347.

61. Sokolowski, M. B., 2001 Drosophila: genetics meets behaviour. Nat. Rev. Genet. 2: 879–890.

62. Spieth, H. T., 1952 Mating behavior within the genus Drosophila (Diptera). Bulletin of the AMNH; v. 99, article 7. Bulletin of the American Museum of Natural History 99: 399–474.

63. Sturtevant, A. H., 1915 Experiments on sex recognition and the problem of sexual selection in Drosophila. Journal of Animal Behavior 5: 351–365.

64. Szafer-Glusman, E., M. G. Giansanti, R. Nishihama, B. Bolival Jr, J. Pringle et al., 2008 A role for very-long-chain fatty acids in furrow ingression during cytokinesis in Drosophila spermatocytes. Curr Biol 18: 1426–1431.

65. Tanji, T., A. Ohashi-Kobayashi, and S. Natori, 2006 Participation of a galactose-specific C-type lectin in Drosophila immunity. Biochem. J. 396: 127–138.

66. Theopold, U., M. Rissler, M. Fabbri, O. Schmidt, and S. Natori, 1999 Insect glycobiology: a lectin multigene family in Drosophila melanogaster. Biochem. Biophys. Res. Commun. 261: 923–927.

67. van de Geijn, B., G. McVicker, Y. Gilad, and J. K. Pritchard, 2015 WASP: allele-specific software for robust molecular quantitative trait locus discovery. Nat Meth 12: 1061–1063.

68. Watanabe, T. K., 1979 A gene that rescues the lethal hybrids between *Drosophila melanogaster* and *D. simulans*. Jpn. J. Genet. 54: 325–331.

69. Wicker-Thomas, C., D. Garrido, G. Bontonou, L. Napal, N. Mazuras et al., 2015 Flexible origin of hydrocarbon/pheromone precursors in Drosophila melanogaster. J. Lipid Res. 56: 2094–2101.

